# Epigenetic clock and methylation studies in gray short-tailed opossums

**DOI:** 10.1101/2021.10.13.464301

**Authors:** Steve Horvath, Amin Haghani, Joseph A. Zoller, Ken Raj, Ishani Sinha, Annais Talbot, Yadiamaris Aviles Ruiz, Karen E. Sears

**Author notes:** Joint first author (Steve Horvath, Amin Haghani). <u>Corresponding authors</u> and. <u>Emails:</u>.

## Abstract

The opossum (*Monodelphis domestica*), with its sequenced genome, ease of laboratory care and experimental manipulation, and unique biology, is the most used laboratory marsupial. Using the mammalian methylation array, we generated DNA methylation data from n=100 opossum tissues including blood, liver, and tail. We contrast age-related changes in the opossum methylome to those of C57BL/6J mice. We present several epigenetic clocks for opossums that are distinguished by their compatibility with tissue type (pan-tissue and blood clock) and species (opossum and human). Two dual-species human-opossum pan-tissue clocks accurately measure chronological age and relative age, respectively. These human-opossum epigenetic clocks are expected to provide a significant boost to the attractiveness of opossum as a biological model.

## INTRODUCTION

*Monodelphis domestica* (the gray short-tailed opossum) is a South American marsupial that is used as a mammalian model system [1–3] in countries including the USA, the UK, Canada, Germany, Poland, Italy, and Australia. Opossums are used as model systems because of their suitability to lab use and the advantages presented by their unique biology relative to placental mammals such as mice. Regarding the former, opossums are small, relatively docile, amenable to laboratory rearing, and have had their genome sequenced [1, 2, 4, 5]. Regarding the latter, opossums are born after only 14-15 days gestation at a developmental stage equivalent to an E10.5-12.5 mouse or a human at 40 days of gestation [1, 6]. Because of their early birth, much of opossum development happens outside the uterus, and can be studied and manipulated more easily than in traditional mammalian models such as mice [1]. This is especially the case as gray short-tailed opossums, although marsupials, lack a pouch. Researchers have taken advantage of these traits to study development of the lung [7], neural crest [8], brain [9–11], limb [12, 13], reproductive [14, 15], and other systems. Because of their early developmental state at birth, newborn opossums are also capable of totally regenerating their spinal cord even after it has been completely severed, and therefore provide an ideal model for spinal cord regeneration [9, 16–18]. Opossums are also one of the only mammals besides humans that develop UV-induced melanoma, making them a good model for melanoma research [19–21]. Opossums are also being used to study many other biomedically-relevant topics, including diet-induced hyperlipidemia [22, 23], wound healing [24], teratogenesis [25, 26], and susceptibility to infection [27, 28]. Another aspect of marsupial biology that differs from that of placentals is their longevity; marsupials, on average, live only about 80% as long as similarly-sized, non-flying placentals [29]. Beyond biomedical topics, opossums have been used to study a suite of evolutionary topics, most of which have compared evolution in marsupial and placental mammals [30, 31].

DNA methylation-based age estimators have been developed for many mammalian species but not yet for opossums [32–40]. Here, we present highly accurate epigenetic clocks that apply to both humans and opossum. The human-opossum clock increases the probability that findings in opossums will translate to humans, and vice versa. Since opossums are already being used in biomedical research, we expect that these clocks will facilitate research into development and the influence of age on pathology.

## Methods

### Opossum and mouse samples

Opossum samples come from a pedigreed, breeding colony of gray short-tailed opossums (*Monodelphis domestica*) that was established by founder individuals purchased from the Southwest Foundation for Biomedical Research. Mouse samples come from a breeding colony of C57BL/6J mice originally purchased from the Jackson Laboratory. Both colonies are maintained by the Sears Lab at UCLA. Opossums were euthanized by CO_2_ inhalation to effect followed by bilateral thoracotomy. Mice were euthanized by CO_2_ inhalation to effect followed by cervical dislocation. These procedures are in accordance with the AVMA Guidelines for the Euthanasia of Animals 2013: https://www.avma.org/KB/Policies/Documents/euthanasia.pdf, and all animal procedures were approved by the UCLA IACUC. All tissue samples, e.g., liver, blood, tail (see Table 1), were taken from euthanized opossums and mice and stored at −20° C until use. DNA from samples was extracted and purified using a DNA Miniprep Plus Kit (Zymo), following manufacturer’s protocols. PicoGreen fluorescent dsDNA was used to assess the concentration of resulting DNA and concentrations adjusted to 50 to 250 ng/ul. DNA samples were submitted to the Technology Center for Genomics & Bioinformatics at UCLA for generation of DNA methylation data and further analyses.

**Table 1.** Description of the opossum and mouse methylation data. Tissue type, N=Total number of samples/arrays. Number of females. Age: mean, minimum and maximum.

| **Species** | **Tissue** | **N** | **No.**<br>**Female** | **Mean**<br>**Age** | **Min.**<br>**Age** | **Max.**<br>**Age** |
| --- | --- | --- | --- | --- | --- | --- |
| Opossum | Ear | 45 | 28 | 0.771 | 0.0384 | 2.22 |
|  | Liver | 48 | 26 | 0.922 | 0.0384 | 3.27 |
|  | Tail | 7 | 4 | 0.164 | 0.0384 | 0.288 |
| Mouse | Blood | 10 | 1 | 0.0595 | 0.0192 | 0.115 |
|  | Ear | 18 | 1 | 0.0671 | 0.0192 | 0.115 |
|  | Liver | 23 | 1 | 0.0684 | 0.0192 | 0.115 |
|  | Muscle | 18 | 1 | 0.0671 | 0.0192 | 0.115 |
|  | Tail | 18 | 1 | 0.0671 | 0.0192 | 0.115 |
|  | Whole<br>Brain | 18 | 1 | 0.0671 | 0.0192 | 0.115 |

### Human tissue samples

To build the human-opossum clock, we analyzed previously generated methylation data from n=1211 human tissue samples (adipose, blood, bone marrow, dermis, epidermis, heart, keratinocytes, fibroblasts, kidney, liver, lung, lymph node, muscle, pituitary, skin, spleen) from individuals whose ages ranged from 0 to 93. The tissue samples came from several sources. Tissue and organ samples from the National NeuroAIDS Tissue Consortium [41]. Blood samples from PEG and the Cape Town Adolescent Antiretroviral Cohort study [42]. Skin and other primary cells provided by Kenneth Raj [43]. Ethics approval (IRB#15-001454, IRB#16-000471, IRB#18-000315, IRB#16-002028).

### DNA methylation data

To overcome the species barrier, we used a DNA methylation array platform (HorvathMammalMethylChip40) that encompasses CpGs flanked by DNA sequences that are conserved across different species of the mammalian class [44]. All methylation data were generated using the mammalian array platform. Not all CpGs on the array apply to opossums. The particular subset of species for each probe is provided in the chip manifest file at the NCBI Gene Expression Omnibus (GEO) platform (GPL28271). The SeSaMe normalization method was used to define beta values for each probe [45].

### Relative age estimation

To introduce biological meaning into age estimates of opossums and humans that have very different lifespan; as well as to overcome the inevitable skewing due to unequal distribution of data points from opossums and humans across age range, relative age estimation was made using the formula: Relative age= Age/maxLifespan where the maximum lifespan for the two species was chosen from an updated version of the *anAge* data base [46]. According to the “anAge” data base [46, 47], the maximum lifespan of Monodelphis domestica is 4.2 years.

### Clocks and penalized regression

Details on the clocks (CpGs, genome coordinates) and R software code are provided in the Supplement. Penalized regression models were created with glmnet [48]. We investigated models produced by elastic net” regression (alpha=0.5 in glmnet R function). The optimal penalty parameters in all cases were determined automatically by using a 10 fold internal cross-validation (cv.glmnet) on the training set. By definition, the alpha value for the elastic net regression was set to 0.5 (midpoint between Ridge and Lasso type regression) and was not optimized for model performance. We performed a cross-validation scheme for arriving at unbiased (or at least less biased) estimates of the accuracy of the different DNAm based age estimators. One type consisted of leaving out a single sample (LOOCV) from the regression, predicting an age for that sample, and iterating over all samples. We subset the set of CpG probes to those that uniquely mapped to a CpG site in the swine genome. While no transformation was used for the blood clock for opossums, we did use a log linear transformation for the dual species clock of chronological age (Supplement). The accuracy of the resulting clocks was assessed via cross validation estimates of 1) the correlation R between the predicted epigenetic age and the actual (chronological) age of the animal, 2) the median absolute difference between DNAm age and chronological age (mae).

### GREAT analysis

We analyzed gene set enrichments using GREAT [49]. The GREAT enrichment analysis automatically conditioned out (removed) any bias resulting from the design of the mammalian array by using a background set of CpGs that map to horses and are located on the mammalian array. The GREAT software performs both a binomial test (over genomic regions) and a hypergeometric test over genes.

We performed the enrichment based on default settings (Proximal: 50.0 kb upstream, 1.0 kb downstream, plus Distal: up to 1,000 kb) for gene sets implemented in GREAT. To avoid large numbers of multiple comparisons, we restricted the analysis to the gene sets with between 10 and 3,000 genes. We report nominal P values and two adjustments for multiple comparisons: Bonferroni correction and the Benjamini-Hochberg false discovery rate.

### EWAS and Functional Enrichment

EWAS was performed in each tissue separately (bladder, blood, cerebral cortex, kidney, liver, lung) using the R function “standardScreeningNumericTrait” from the “WGCNA” R package [50]. Next the results were combined across tissues using Stouffer’s meta-analysis method. The functional enrichment analysis was done using the genomic region of enrichment annotation tool [49]. CpGs implicated by our EWAS were filtered for CpG position information, lifted over to the human genome using UCSC’s Liftover tool and fed into the online functional analysis tool GREAT using the default mode, to obtain a list of significantly enriched functions for both positive and negative EWAS hits in the different tissues.

## Results

In total, we analyzed n=100 opossum tissue samples from ear, liver, tail as detailed in **Table 1**. Since we aimed to contrast development between opossums and mice, we used the same methylation array platform to profile n=105 mouse tissue samples (blood, liver, tail, ear, muscle, whole brain) during development (postnatal weeks 1,2,3,4,5,6) as detailed in **Table 1**. Unsupervised hierarchical clustering revealed that the opossum and mouse samples clustered by tissue type (**Supplementary Figure S1, Supplementary Figure S2**). To assess whether platemap errors occurred, we constructed random forest predictors to tissue type and sex in each species. These classifiers exhibited perfect accuracy, with respective (out-of-bag) error rates of zero.

### Predictive Accuracy of the Epigenetic Clock

To arrive at unbiased estimates of the epigenetic clocks, we applied cross-validation analysis with the training data. For the development of the basic opossum clock, this consisted of opossum blood, liver, tail DNA methylation profiles. For the generation of human-opossum clocks however, the training data was constituted by human and opossum DNA methylation profiles. Cross-validation analysis reports unbiased estimates of the age correlation R (defined as Pearson correlation between the age estimate (DNAm age) and chronological age) as well as the median absolute error.

From these analyses, we developed three epigenetic clocks for opossums that vary with regards to two features: species and measure of age. The pan-tissue opossum clock was trained on 3 tissues (ear, liver, tail) and is expected to generalize to other tissues as well.

The two human-opossum clocks mutually differ by way of age measurement. One estimates chronological ages of opossums and humans (in units of years), while the other estimates relative age, which is the ratio of chronological age of an animal to the maximum lifespan of its species; with resulting values between 0 and 1. We prefer the human opossum clock for relative age since it turns out to be more accurate and because it is arguably more meaningful for biological studies. The relative age estimate facilitates a biologically meaningful comparison between species with very different lifespans such as opossum (4.2 years) and human (122.5 years), which cannot otherwise be afforded by direct comparison of their chronological ages.

As indicated by its name, the pure opossum clock is highly accurate in age estimation of all the tested opossum tissues (R=0.85, and median error=0.32 years, **Figure 1a**). The pan-tissue clocks exhibit high age correlations in individual opossum tissues (R>=0.85, **Supplementary Figure S3**). While the pure opossum clock exhibits a very high correlation with age this does not imply high concordance with chronological age: most opossum samples were estimated to have ages that are considerably different from their chronological age as reflected by a relatively high median absolute error.

**Figure 1.**
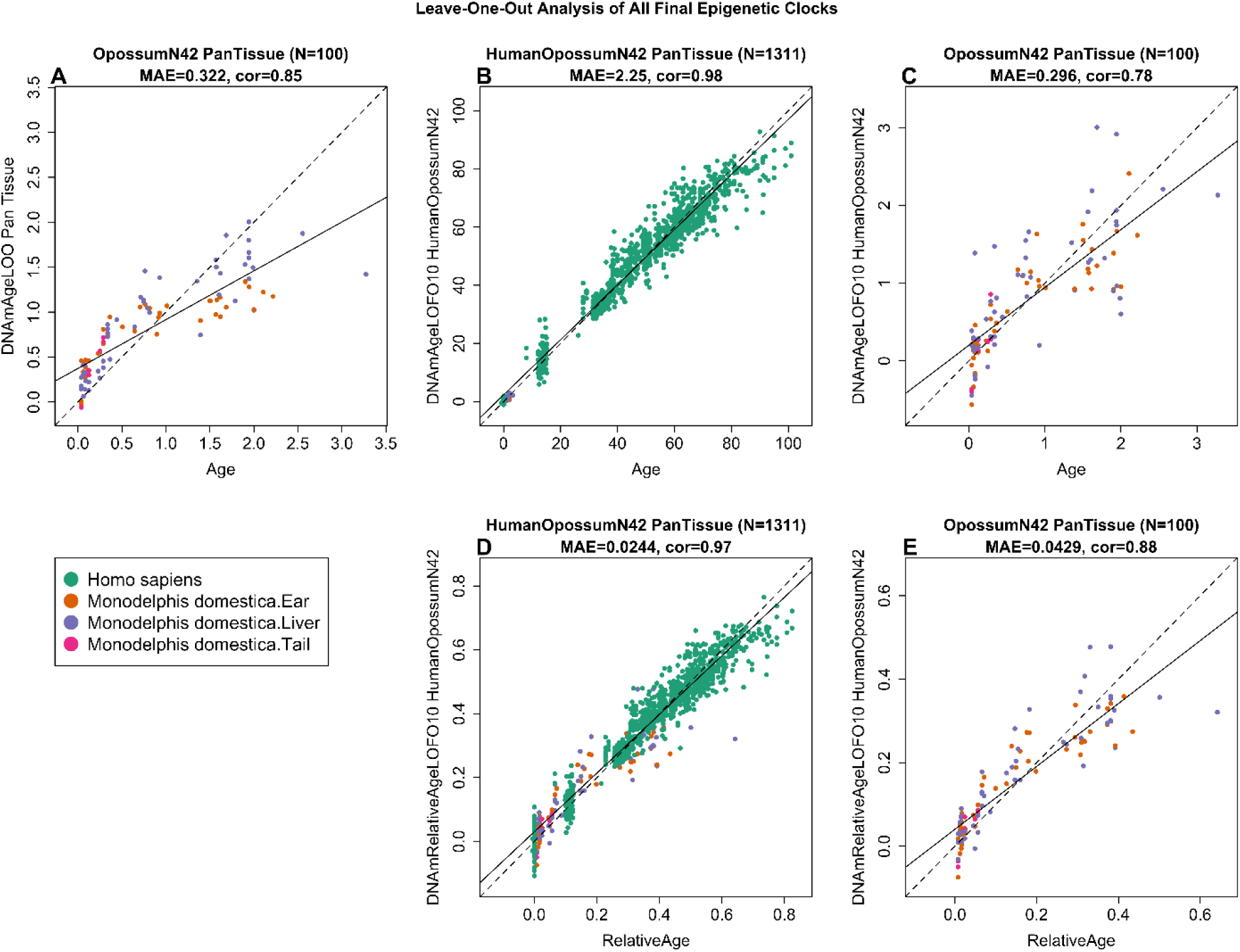
Cross-validation study of 3 epigenetic clocks for opossums. We developed 3 epigenetic clocks for opossums: A) pan-tissue clock that was trained with opossum tissues only, B,C) human-opossum clock for chronological age applied to B) both species and C) opossums only. Human-opossum clock for relative age applied to D) both species and E) opossums only. Leave-one-sample-out (LOO) estimate (y-axis, in units of years) versus chronological age or relative age (x-axis). Relative age is defined as chronological age divided by the maximum age of the respective species. The linear regression of epigenetic age is indicated by a solid line while y=x is depicted by a dashed line.

The human-opossum clock for chronological age leads to high age correlations when DNA methylation profiles of both species are analyzed together (R=0.98, **Figure 1b**), but is less accurate when restricted to opossum tissue samples (R=0.78, **Figure 1c**). The human-opossum clock for relative age exhibits superior accuracy when restricted to opossums (R=0.88, **Figure 1e**). The use of relative age circumvents the clustering of data points of opossums and humans to opposite parts of the curve, which is evident in Figure 1b. The human-opossum clock of relative age is particularly exciting as it allows comparison between humans and opossums based on their relative positions within the lifespans of both species.

### EWAS of chronological age

The mammalian methylation array contains 15,098 CpGs that map to *Monodelphis domestica* (ASM229v1.100) genome. Due to the high inter-species conservation of the probes on the array, findings from the opossum methylation data can probably be extrapolated to humans and other mammalian species. We performed an epigenome-wide association analysis (EWAS) of age for these CpGs in ear, liver, and tail of opossums. The EWAS analysis in tails was arguably underpowered due to low sample size (n=7). At a nominal significance of p<10^−4^, a total of 2971, 1328, and 107 CpGs were related to age in ear, liver, and tail of opossums, respectively (**Figure 2a**). The top age-associated CpGs and their proximal genes for the individual tissues are as follows: ear, a decrease of methylation in ENSMODG00000019303 promoter (p=2×10^−18^), and MSTN exon (p=8×10^−17^); liver, a decrease of methylation in SRSF5 exon (p=5×10^−18^); and tail, an increase of methylation in *PSMD7* intron (p=1×10^−7^). In general, these CpGs were enriched with genes related to RNA processing (p=8×10^−15^), RNA splicing (3×10^−14^), cell cycle (3×10^−18^), and replicative senescence (1×10^−7^) (**Figure S4**).

**Figure 2.**
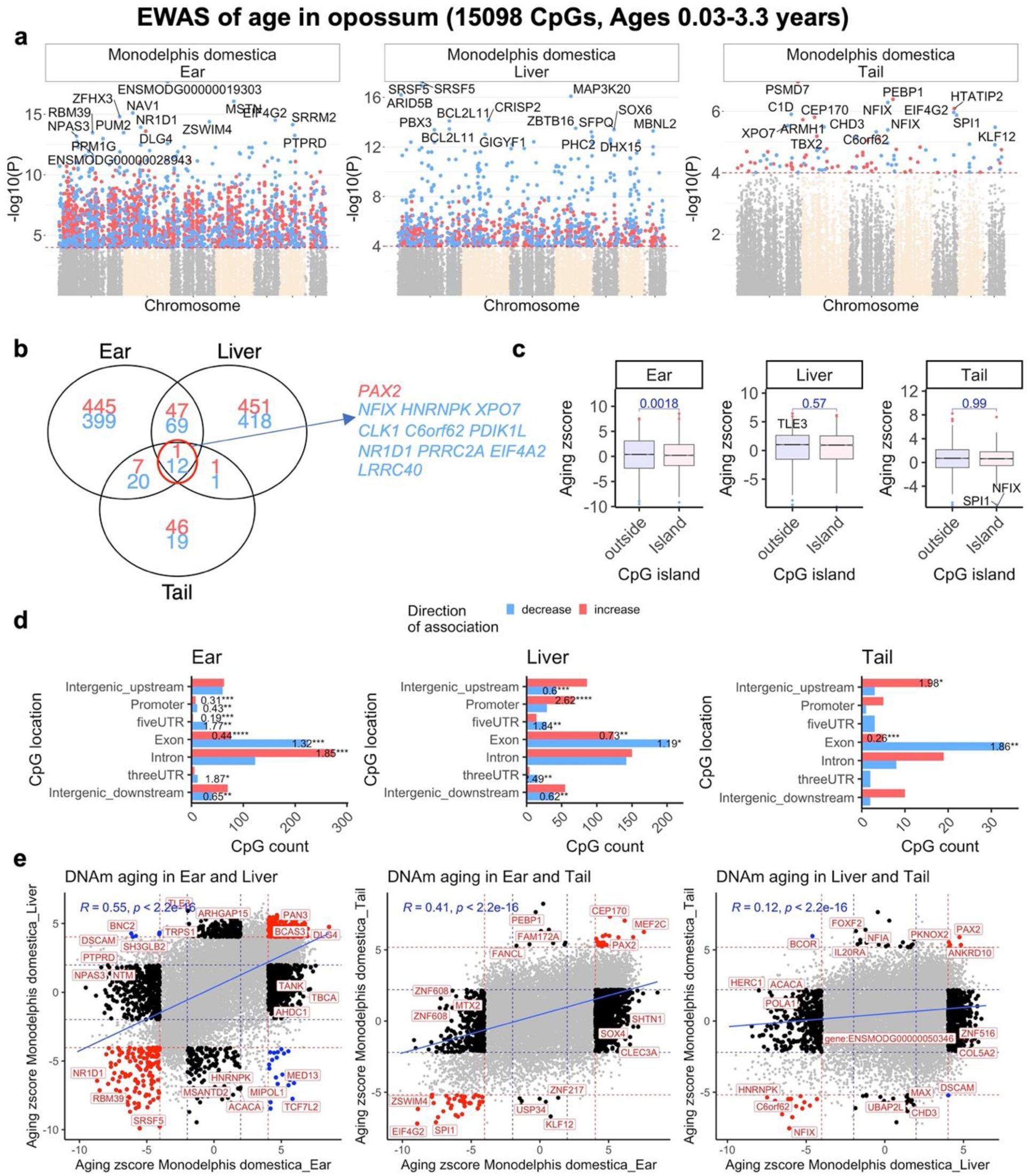
Epigenome-wide association (EWAS) of chronological age in opossum tissues. EWAS of age in ear (n=45), liver (n=48), and tail (n=7) of opossum. **a**, Manhattan plots of the EWAS of chronological age. The coordinates are estimated based on the alignment of Mammalian array probes to Monodelphis_domestica.ASM229v1.100 genome assembly. The direction of associations with p<10^−4^(red dotted line) is highlighted by red (increase) and blue (decrease) colors. Top 15 CpGs was labeled by the neighboring genes. **b**, Venn diagram of the overlap in up to top 1000 (500 in each direction) significant CpGs in each tissue. **c**, box plots of DNAm aging by CpG island status. The x axis is the Fisher transformed z score of the Pearson correlation of each CpG with age. The t-test p values are labeled above the box plots. **d**, Location of top CpGs in each tissue relative to the closest transcriptional start site. The odds ratio of the proportion changes than the background are reported in for each bar. Fisher exact p values: * p<0.05, ** p<0.01, ***p<0.001, ****p<0.0001. **e**, Scatter plots represent the pair wise comparison of EWAS of age in different opossum tissues. The aging z scores are the Fisher z-transformation of DNAm-Age Pearson correlation for each CpG in opossum tissues. A positive (or negative) of z score means an increase (or decrease) of DNAm with age in the analyzed species. Red dotted line are the z scores corresponding to p<10-4; blue dotted line are the z scores corresponding to p>0.05; Red data dots indicate shared CpGs (i.e. The CpGs that significantly change in the same direction in both tissues) between tissue represented by the X and Y axes; black data dots are CpG methylation changes that are significant in one but not the other tissue. The top CpGs in each sector is labeled by the identity of the adjacent genes.

EWAS of chronological age revealed a clear tissue specific DNAm change in opossums, however, these results are confounded by different age ranges of analyzed tissues. While the liver (10 days to 3.2 years) and ear (10 days to 2.2 years) were available from all ranges of opossum lifespan, the tail samples were collected during development (10 days to 0.28 years), and the underlying sample size was very low (n=7, Table 1). Despite these caveats, our analysis identified 13 CpGs with methylation changes that were shared between these three tissues, in the top 1000 age-related CpGs (500 per direction of association) (**Figure 2b; Figure S5**).

The opossum CpG islands did not show a systematic increase of methylation with age (**Figure 2c**), which suggests a unique DNAm aging pattern in marsupials compared to other mammals. This observation was more prominent for ear samples. In opossum ear, the CpG islands had a slightly lower aging association than CpGs outside of islands (t-test p=0.0018) (**Figure 2c**). In addition, there were relatively few methylation changes in CpGs located in promoters (OD=0.31, Fisher Exact p<10^−3^) and 5’UTRs (OD=0.19, Fisher Exact p<10^−3^) of genes (**Figure 2d**). These features contrast with placentals, which generally experience age-related gain of methylation in CpG islands, promoters and 5’UTRs [51]. To further compare the tissues, we created pairwise sector plots of DNAm aging in opossum samples. We observed a high positive correlation of age-related DNAm changes between ear-liver (r=0.55), and ear-tail samples (r=0.45) (**Figure 2e**). In contrast to these similarities, there were some age-related CpGs with methylation changes that were divergent between opossum tissues, e.g. *BNC2* (intron) shows a methylation increase in liver but a decrease in ear. This finding demonstrates that aging patterns in one tissue typically differ from those of another tissue.

### Comparing developmental DNAm aging between opossum and mouse tissues

As described above, opossum DNAm aging displays some unique aspects such as a lower degree of change in CpG islands than placental mammals. This observation encouraged us to do a direct comparison of opossum and mouse tissues. We limited the analysis to the developmental stages of these two animals (ages < 1.6 months). Moreover, only a subset of 8819 CpGs that could be mapped to orthologous genes in both species were included in the analysis. At p<0.005 (FDR<3×10^−6^ to <0.1 depending on the species-tissue), mouse tissues (average change = 2632 CpGs) had a higher number CpGs related to age than opossum tissues (average change = 488) (**Figure 3a**). This difference can have a biological origin as we have a comparable sample size, particularly in ears, for these two species (16 opossum ears, 18 mouse ears). Despite this difference, there were nevertheless a considerable number of CpGs that experienced similar methylation changes during development of the mouse and opossum in all three tissues (mean correlation = 0.2) (**Figure 3b**). Interestingly, 3-10% of the top 1000 age-related CpGs in mouse were shared with opossum tissues (**Figure 3c**). This observation shows that the development process is partially conserved between these two species. The shared aging patterns were enriched with genes related to development (p = 6×10^−10^), polycomb repressor complex 2 binding sites (p = 4×10^−8^), and H3K27ME3 marks (p=2×10^−8^), which are a common aging pattern in all previously analyzed placental species [51]. Polycomb family protein complex maintains the transcriptional repressive states of genes. These proteins also regulate H3K27Me3 marks, DNA damage, and senescence states of the cells during aging [52]. We also identified 8 to 38 CpGs with a divergent aging pattern in mouse and opossum depending on the tissue (**Figure S6**). Interestingly, these divergent CpGs were adjacent to genes involved with the immune system (e.g., lymphocyte differentiation, p=5×10^−4^) and the female reproductive system (e.g., luteinization, p = 4×10^−5^) (**Figure 4; Figure S7** shows the top terms for each species-tissue strata).

**Figure 3.**
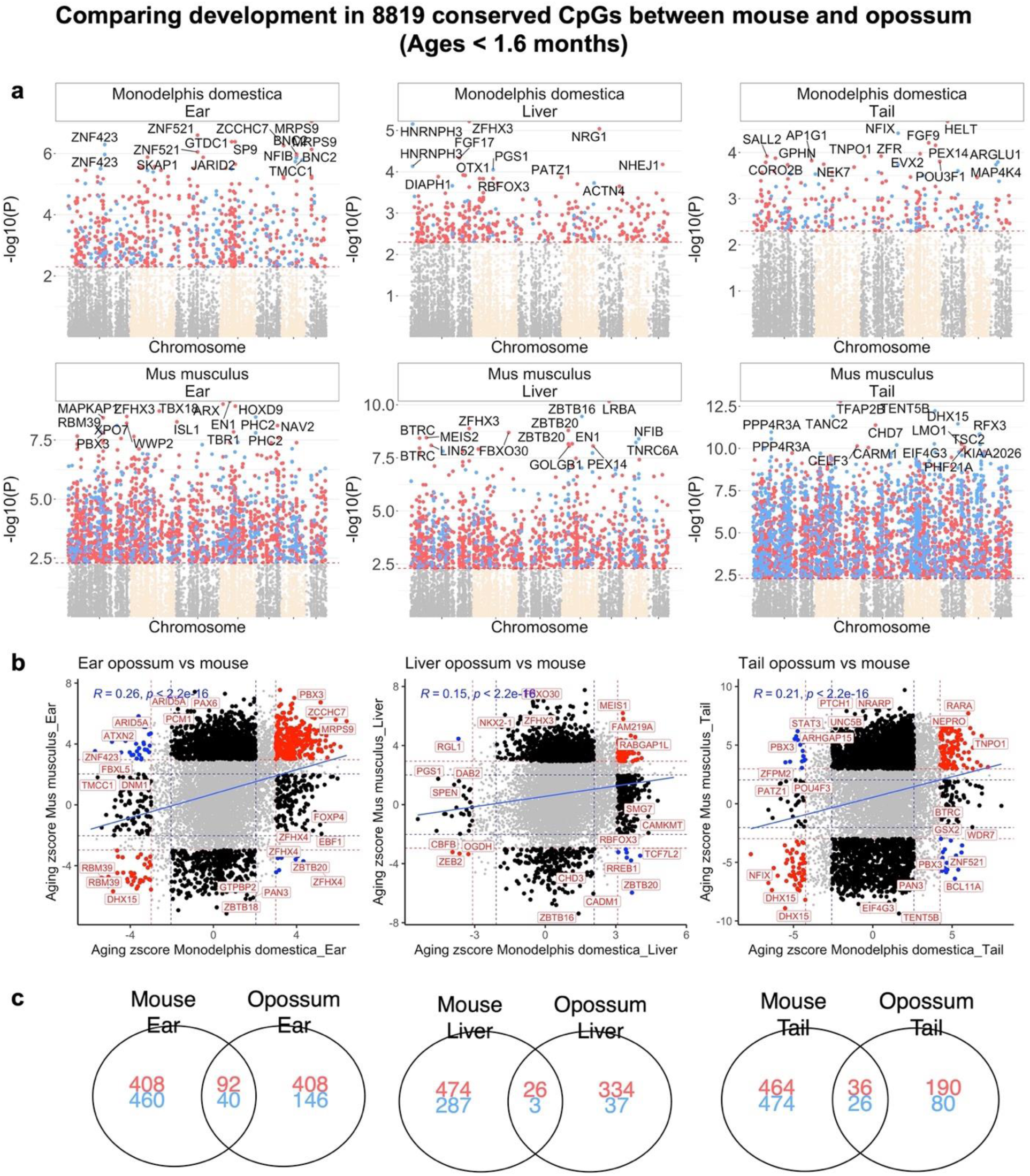
Comparing DNAm aging between mouse and opossum. **a**, Manhattan plots of the EWAS of chronological age in developmental stages of mouse and opossum. The analysis is limited to 8819 conserved CpGs in these two species. All coordinates are reported based on Monodelphis_domestica.ASM229v1.100 genome assembly. The direction of associations with p<0.005 (red dotted line) is highlighted by red (increased methylation) and blue (decreased methylation) colors. Top 15 CpGs was labeled by the neighboring genes based on the opossum genome. **b**, Scatter plots represent the pair wise comparison of EWAS of age in mouse and opossum. The aging z scores are the Fisher z-transformation of DNAm-Age Pearson correlation for each CpG in different tissues. A positive (or negative) z scores means an increase (or decrease) methylation with age in the analyzed species-tissue strata. Red dotted line are the z scores corresponding to p<0.005; blue dotted line are the z scores corresponding to p>0.05; Red dots indicates shared CpGs with methylation that change significantly in the same direction in both tissues represented by X and Y axes; black dots represent CpGs with methylation that change in one species but not the other. Gene adjacent to the top CpGs in each sector is labeled by opossum gene symbol. **c**, Venn diagram of the overlap of up to 1000 (500 in each direction) significant CpGs between mouse and opossum for each tissue. It is noteworthy that there is so little similarity between the livers of mouse and opossum, which probably reflects the substantial heterogeneity in the liver samples from opossums (Figure S1). On the other hand, much more similarities are seen for tails of these animals, even though the number of opossum tail samples was very small (n=7), which would intuitively lead one to expect lower likelihood of detecting similarities.

**Figure 4.**
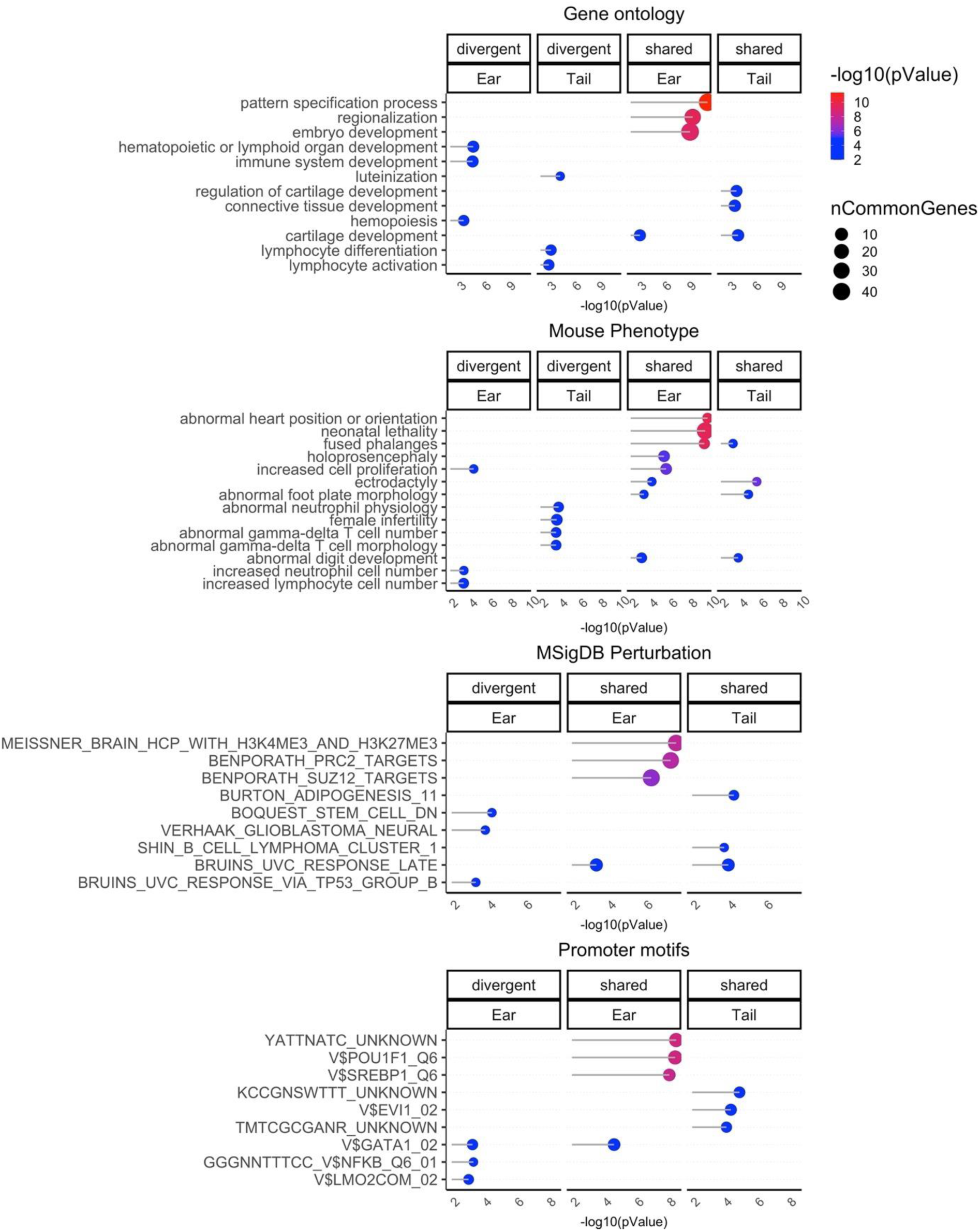
Gene set enrichment analysis of DNAm aging in mouse and opossum. The gene level enrichment was done using GREAT analysis [49] and human Hg19 background limited to 8819 conserved CpGs between mouse and opossum. The CpGs are annotated with adjacent genes within 50kb. We extracted up to top 500 CpGs based on p value of association per direction of change as input for the enrichment analysis. The p values are calculated by hypergeometric test of the EWAS results with the genes in each background dataset. Datasets: gene ontology, mouse phenotypes, promoter motifs, and MsigDB Perturbation, which includes the expression signatures of genetic perturbations curated in GSEA database. The results were filtered for significance at p < 10^−3^ and only the top terms for each EWAS are reported. The shared set means a similar age-related change in both species. The divergent group indicates the CpGs with an opposite age-related change between mouse and opossum.

## Discussion

We previously developed dual species clocks for different placental species [32–40]. A critical step toward building dual species clocks was the development and use of a mammalian DNA methylation array that profiles CpGs with flanking DNA sequences that are highly conserved across numerous mammalian species. The employment of the mammalian array to profile 100 opossum tissues from 3 opossum tissues allowed us to construct a pan-tissue opossum clock, which will be very useful in research that employs opossum as the animal model of choice. The utility of such an epigenetic clock can be further enhanced if its “read-out” can be correlated directly to human age. This would ease the translation of findings from the opossum to humans and *vice versa*. This rationale, which drove us to create dual species clocks with numerous other animals, led us to develop human-opossum clocks. Towards this end, in addition to DNA sequence differences between humans and opossums, we had to also contend with markedly different lifespans between these two species. The maximum lifespan of *Monodelphis domestica* is only 4.2 years according to the “anAge” data base [46, 47], which is 29 times shorter than that of humans (maximum lifespan 122.5). This is an important consideration because the biological fitness of a 3-year-old opossum is not equivalent to that of a 3-year-old human. This challenge is addressed by the introduction of the concept of relative age. An opossum with a relative epigenetic age of 0.5 is much more comparable to a human of the same relative age. The human-opossum clock for relative age can be applied to both humans and opossum-based models of diseases and health conditions. This will encourage investigations into the epigenetic correlates of development and aging. These clocks will also allow evaluation of the impact of environment, living condition, food, and treatment on the rate of opossum epigenetic aging, which are readily translated to humans using the dual species clocks.

While the CpGs that constitute the clocks are the most accurate with regards to age estimation, they are not the only CpGs with methylation changes that alter with age. Indeed, the clock CpGs are only a small subset of all the age-related CpGs. Hence efforts to accrue biological meaning of age-related methylation changes necessitate that all age-related CpGs are analyzed in an epigenome-wide association study (EWAS). Such analyses uncovered a few unexpected features, of which the absence of increased methylation of CpG island with age in opossums is particularly interesting as all placental species studied thus far exhibit this feature. This difference may be linked to another which is that opossum genome possesses a very low overall level of CpG content; on the order of half or less the content of other amniotes [5, 53, 54]. Marsupial genomes are generally of comparable size with their placental counterparts, but are typically packaged into fewer, very large chromosomes [5]. Marsupials’ low CpG content has been hypothesized to be due to the generally lower rate of recombination in marsupials in general [53]. As such, compared to most other mammalian species, opossums have a low density of CpG islands (7.5 per Mb) and a few large chromosomes; consistent with the observation that large chromosomes are correlated with low CpG island density [54].

Our analyses revealed a high degree of tissue-specific age-related methylation changes. This is to be expected as methylation is a means by which expression of genes are regulated in different tissues. At this early stage of our understanding of epigenetics, it is not possible to determine from first principles, which CpG methylation impacts gene expression and, if it does, of which genes. Hence we employed a conservative approach of linking CpGs to the potential regulation of genes that are adjacent and proximal to them. This identified distinct sets of potential age-related genes in the ear, liver, and tail. One of the highest scoring potential age-related genes of the ear is MSTN, which encodes myostatin. This is interesting because myostatin level rises with age, with the highest levels in frail individuals [55]. This is connected to myostatin’s association with muscle loss and dysfunction of muscle stem cells [56]. In the tail, the highest scoring age-related CpG is adjacent to the gene *PSMD7*, which is a subunit of the proteasome, which is essential for the destruction of incorrectly folded and dysfunctional proteins. Indeed, reduced efficiency of protein homeostasis is one of the major hallmarks of aging. Furthermore, it was reported that reduced expression of *PSMD7* leads to reduced mTOR level and activity. This is relevant because perturbation of mTOR activity impacts on the rate of epigenetic aging [57]. It is important to note that at this stage of understanding, these connections uncover only potential associations. They do not yet allow us to know the direction of gene expression change and the outcome. The complexity of these changes can be further compounded especially by candidate genes that encode proteins with potentially higher level of impact. As a case in point, in the opossum liver, the highest scoring age-related CpG is proximal to *CRCF5*, which encodes a splicing factor that carries out alternative splicing of precursor mRNAs. Such a candidate protein can induce cellular impacts that are orders of magnitude greater since they affect the expression of many other genes. Elucidation of potential age-related outcome of such epigenetic change is very complex and requires empirical investigations.

Another set of potential age-related proteins with equally complex outcomes are transcription factors. These proteins affect the expression many other genes. Hence a non-empirical deduction of cause to effect is impossible. As a case in point, one of the highest-scoring age-related CpGs that are common to all the three opossum tissues lie next to the *PAX2* gene, which encodes a transcription factor that is a member of PAX protein family. Proteins of this family participates in the regulation of developmental genes across a large swathe of species. Indeed, *PAX2* was also identified as one of the potential age-related genes in EWAS carried out with baboons, while other members of the Pax family are implicated in epigenetic aging of other species. All these highlight the conservation of the developmental process across mammals and the association of development with aging. This very strong link is further consolidated by the implication of the polycomb repressive complex in epigenetic aging. Regardless of species phylogeny and their evolutionary journey, polycomb repressive complex is identified as being associated with aging in all mammalian species analyzed to date, bar none.

It is of further interest that two high-scoring, age-related CpGs that are shared between all three opossum tissues lie adjacent to the *NFIX* gene, which encodes yet another transcription factor. Interestingly, mutation of the *NFIX* gene is associated with Sotos syndrome 2, which is a developmental disorder characterized by excessive growth and advanced bone age. Indeed, Sotos syndrome patients exhibit accelerated epigenetic aging [58].

Collectively, the conservative approach of looking at genes that are proximal to age-related CpGs have uncovered multiple, tantalizing potential age-related genes. It is acknowledged that epigenetic changes can have distal effects too, and when the technology and methodology to analyze these become available, we will have a more comprehensive understanding of epigenetic aging.

Comparison between developing opossum and mouse revealed expected similarities and also interesting differences, of which genes encoding constituents of the immune system are prominent in developing opossum. In contrast to the situation in placentals, much of the development of the adaptive immune system occurs after birth in marsupials [59, 60]. For example, opossum newborns do not begin to produce their own antibodies until a week or more after their birth [61]. As a result, with only their innate immune system up and running, newborn marsupials must navigate the ex-utero world and its potentially pathogenic microorganisms [62]. Survival of the newborn in this relatively unprotected state is thought to depend heavily on antimicrobial proteins obtained through the mother’s milk [63–66]. Consistent with this, the marsupial genome displays expansions in some components of the innate immune system, including the cathelicidin and defensin families of antimicrobial peptides [61, 63].

In conclusion, we present here, the epigenetic clocks for opossum that will be of use and interest to those who employ these developmentally unique creatures as animal models. The close association between development and aging is once again revealed, and the potential age-related genes identified here presents the opportunity to investigate the biology and process of aging in these marsupials, and the human-opossum clocks will be instrumental in translating these finds to human aging.

## Author contributions

SH and KS conceived of the study and wrote the article. AH, JZ, SH carried out the statistical analysis. The remaining authors helped with interpreting the data and data generation. All authors reviewed and edited the article.

## Acknowledgements

This work was supported by the Paul G. Allen Frontiers Group (SH) and by NIH grant R21OD022988-03.

## DATA AVAILABILITY

The data will be made publicly available as part of the data release from the Mammalian Methylation Consortium. Genome annotations of these CpGs can be found on Github https://github.com/shorvath/MammalianMethylationConsortium

The mammalian methylation array is distributed by the Epigenetic Clock Development Foundation.

