## Supplementary Figures and Tables for "Epigenetic clock and methylation studies in gray short-tailed opossums"

**SUPPLEMENTARY MATERIAL**

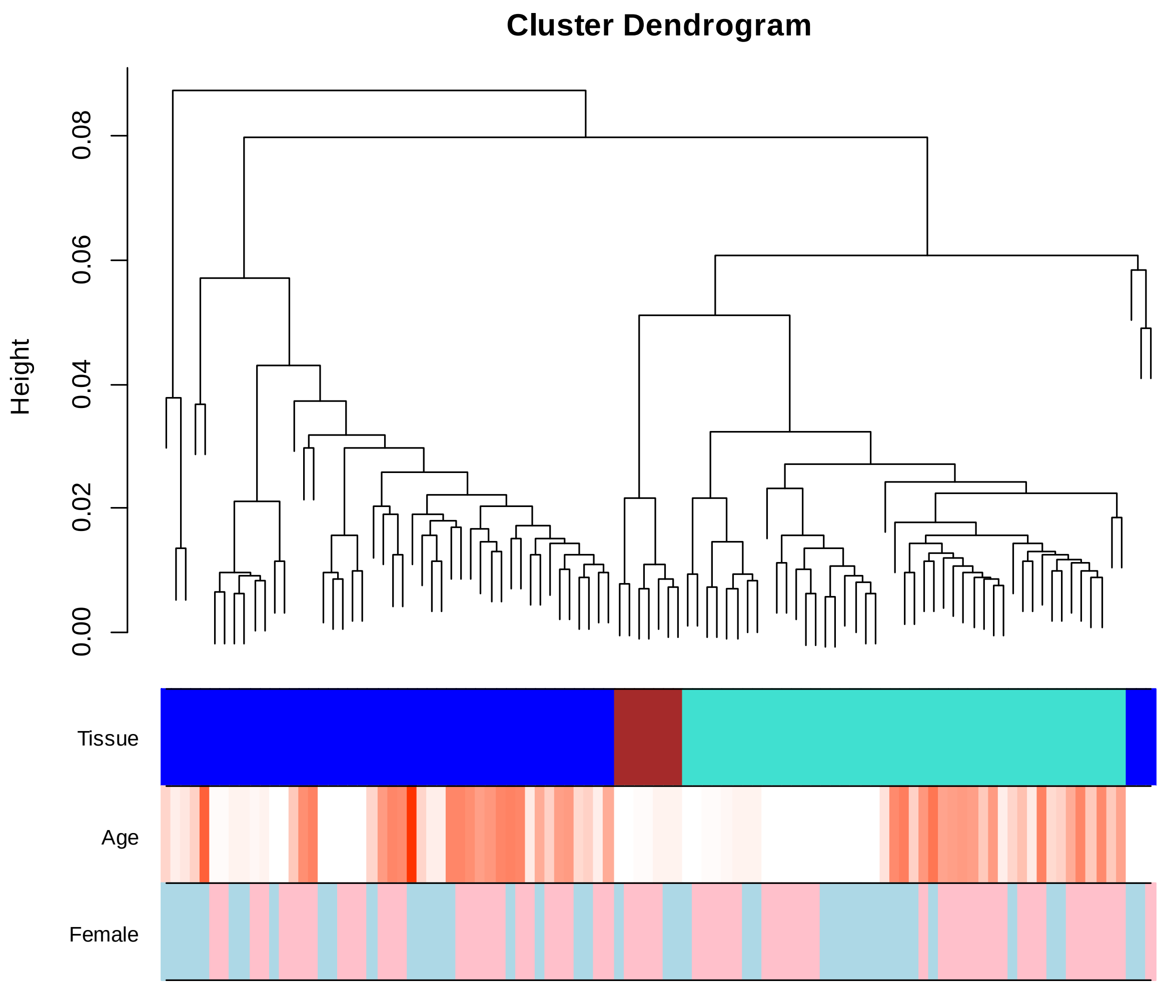


**Figure S1. Unsupervised hierarchical clustering in opossum**. Average linkage hierarchical clustering based on the interarray correlation coefficient (Pearson correlation). The first color-band underneath the tree color codes tissue: ear (turquoise), liver (blue), tail (brown). Age color codes numeric values: old (red) versus young (white). The third color band encodes sex (female= pink, male=lightblue). The opossum tissues largely cluster by tissue type but the liver samples (blue) fall into 3 separate branches (clusters) at a height cut-off (y-axis) of say 0.05. The fact that the liver samples don't cluster together could reflect technical noise (resulting from thawing the entire frozen animal) or biological variability.

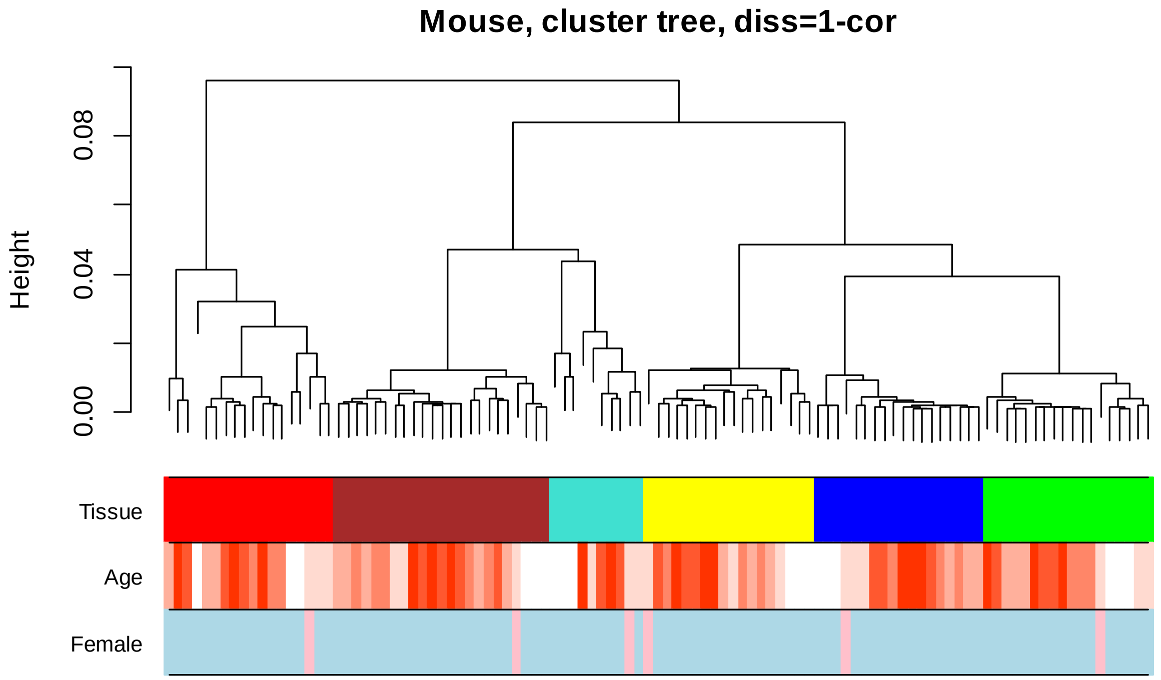


**Figure S2. Unsupervised hierarchical clustering in mice**. Average linkage hierarchical clustering based on the interarray correlation coefficient (Pearson correlation). The first color-band underneath the tree color codes tissue: Blood (turquoise), ear (blue), liver (brown), muscle (yellow), tail (green), whole brain (red). Age color codes old (red) versus young (white). Female= pink, male=lightblue.

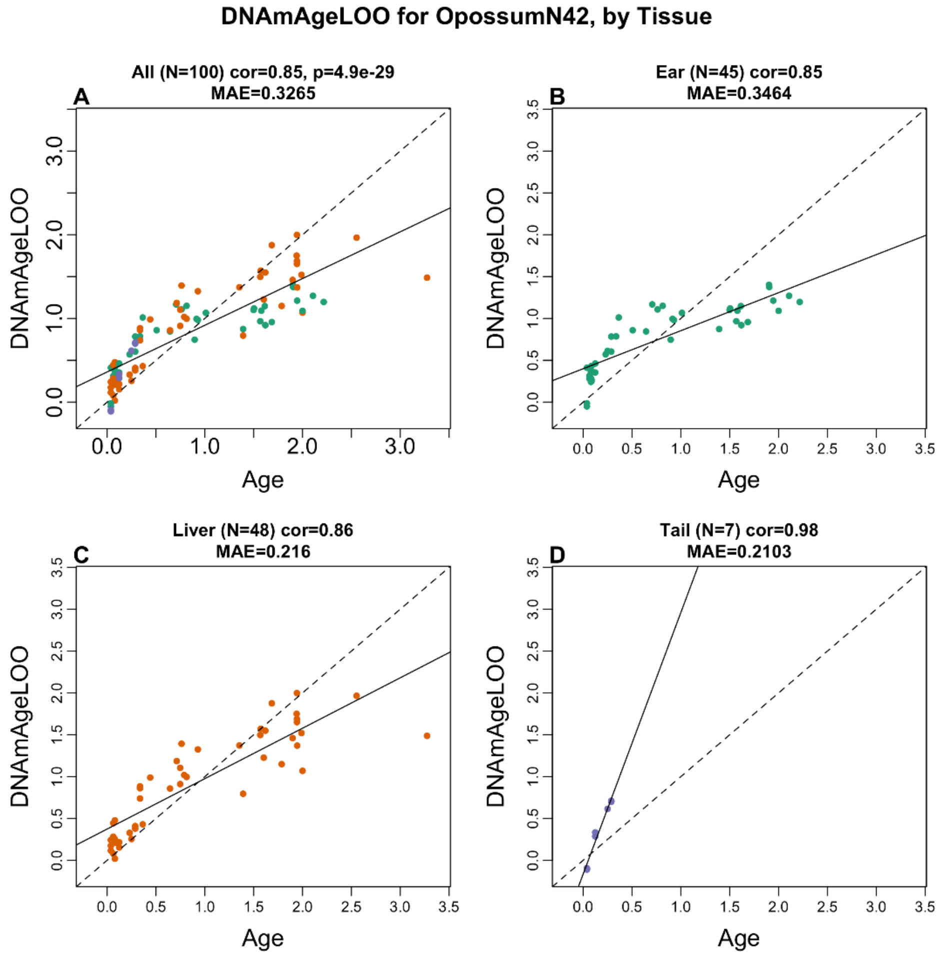


**Figure S3. Pan tissue clock for opossums**. Leave one sample out (LOO) estimates of age (y-axis) based on methylation levels versus chronological age at sample collection (x-axis). All axes are in units of years. A) All tissues combined. Dots (samples) are colored as in the other panels. B) ear samples (green dots),C) liver samples (red dots) ,D) tail samples (blue dots). The title of each panel reports the tissue, the sample size (N), the Pearson correlation coefficient, and the median absolute error.

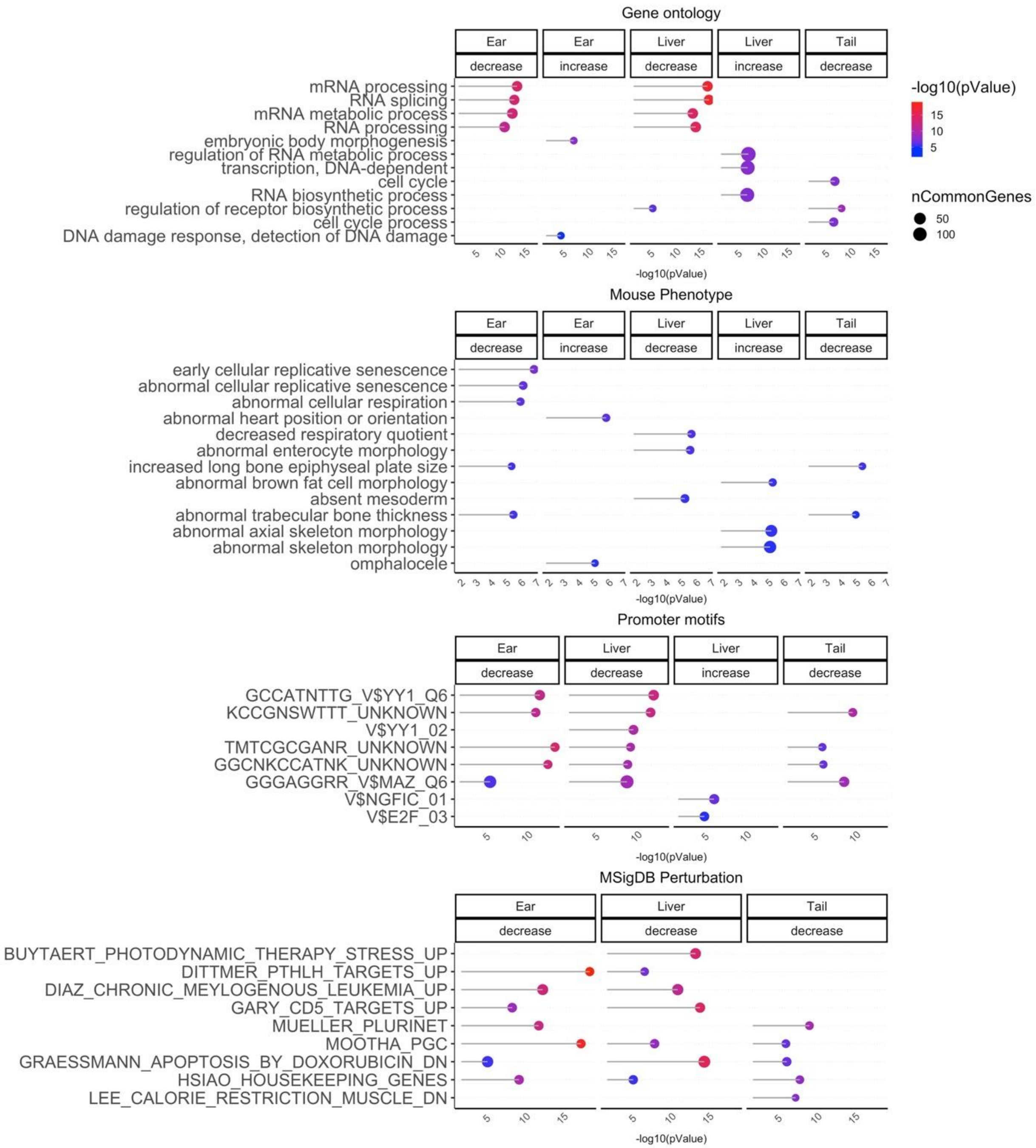


**Figure S4. Gene set enrichment analysis of DNAm aging in opossum tissues**. The gene level enrichment was done using GREAT analysis [49] and human Hg19 background limited to CpGs that could be aligned to opossum. The CpGs were annotate with adjacent genes in 50kb flanking region. We extracted up to top 500 CpGs based on p value of association per direction of change as input for the enrichment analysis. The p values are calculated by hypergeometric test of the EWAS results with the genes in each background dataset. Datasets: gene ontology, mouse phenotypes, promoter motifs, and MsigDB Perturbation, which includes the expression signatures of genetic perturbations curated in GSEA database. The results were filtered for significance at $p < 10^{-3}$ and only the top terms for each EWAS result.

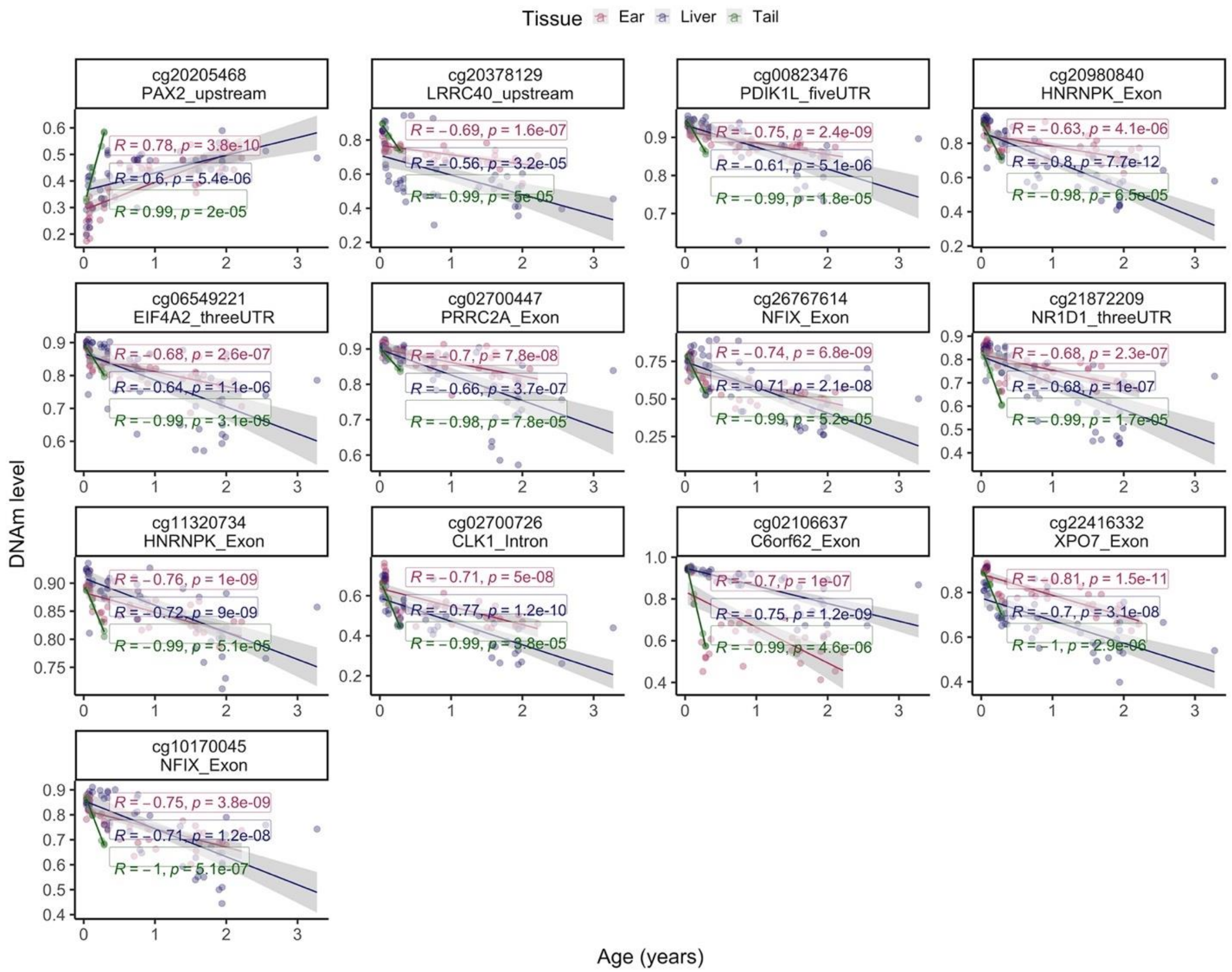


**Figure S5. CpGs with a similar aging pattern in three opossum tissues.** Each panel corresponds to a different CpG with strong age correlations in 3 different tissue types. The methylation values (beta values, y-axis) versus chronological age (in years, x-axis). Each dot corresponds to a DNA sample colored by tissue type: read=ear, green=tail, blue=liver. Each panel reports Pearson correlation coefficients and Student T test p values.

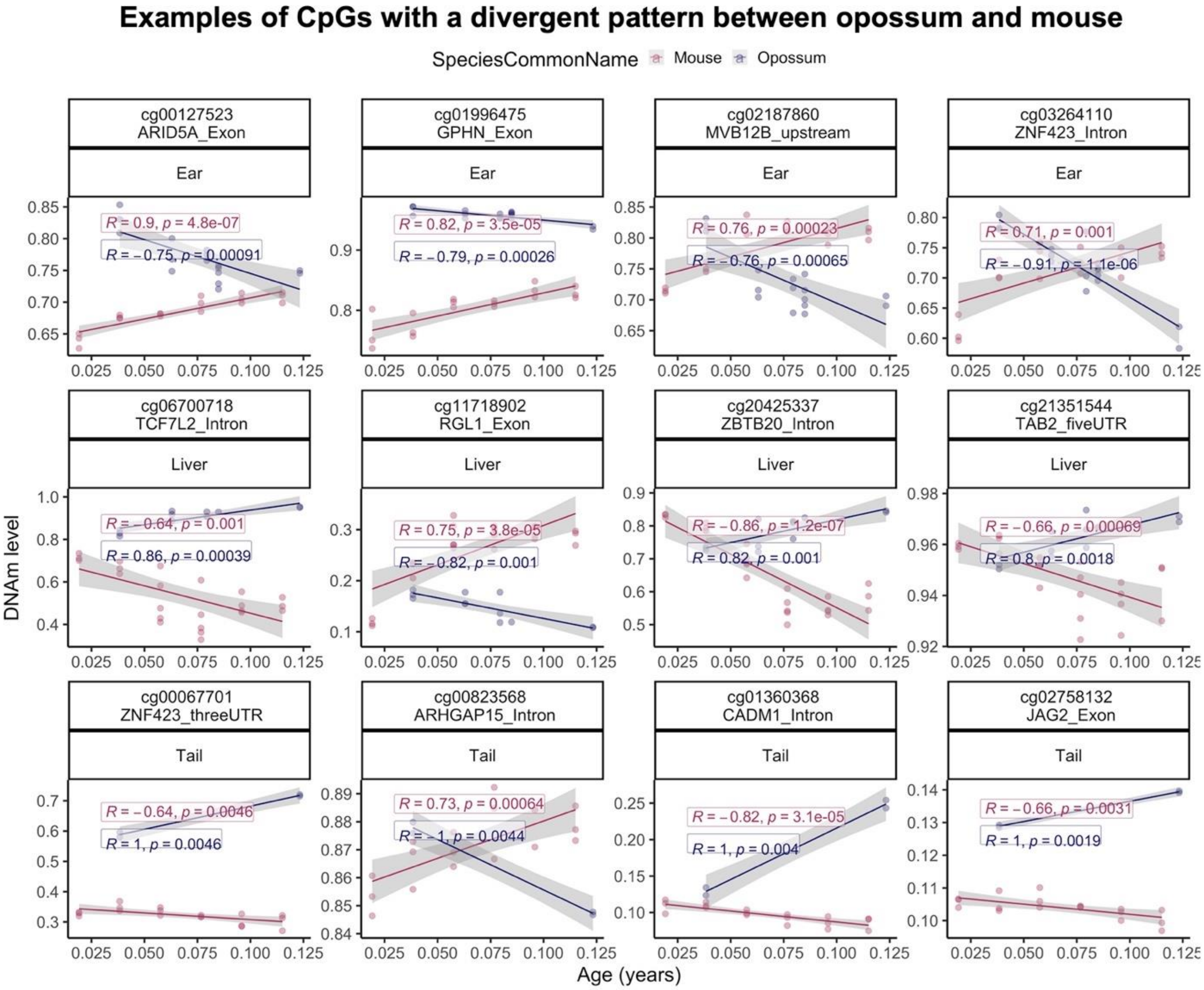


**Figure S6. CpGs whose developmental patterns differ between opossums and mice.** Each panel corresponds to a different CpG whose developmental pattern (aging pattern during development) differs between the two species. The methylation values (beta values, y-axis) versus chronological age (in years, x-axis). Each dot corresponds to a DNA sample colored by species (red=mouse, blue=opossum). The lines correspond to linear regression lines. Each panel reports the tissue type, Pearson correlation coefficients and Student T test p values.

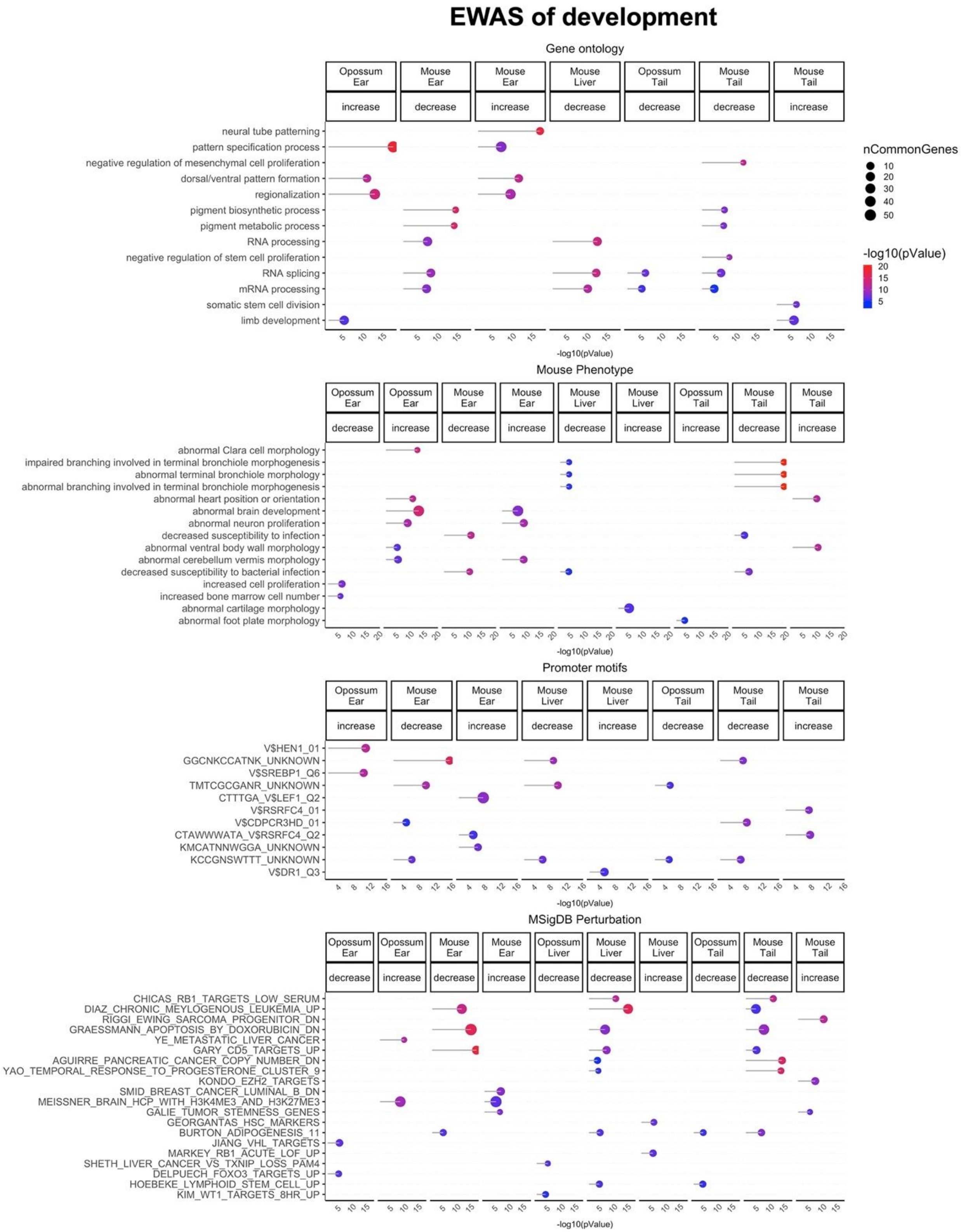


**Figure S7. Gene set enrichment analysis of DNAm aging in opossum tissues**. The gene level enrichment was done using GREAT analysis [49] and human Hg19 background limited to 8819 conserved CpGs between mouse and opossum. The CpGs were annotated by adjacent genes in the 50kb flanking region. As input, we used up to top 500 CpGs based on p-value per direction (age related gain/loss of methylation). The enrichment p-values were calculated using the GREAT analysis. Datasets: gene ontology, mouse phenotypes, promoter motifs, and MsigDB Perturbation, which includes the expression signatures of genetic perturbations curated in GSEA database. Points are colored by significance level as indicated in the legends. All terms are significant at a nominal p value $< 10^{-3}$.
